# Processivity, velocity and universal characteristics of nucleic acid unwinding by helicases

**DOI:** 10.1101/366914

**Authors:** Shaon Chakrabarti, Christopher Jarzynski, D. Thirumalai

## Abstract

Helicases that act as motors and unwind double stranded nucleic acids are broadly classified as either active or passive, depending on whether or not they directly destabilize the double strand. By using this description in a mathematical framework, we derive analytic expressions for the velocity and run-length of a general model of finitely processive helicases. We show that, in contrast to the helicase unwinding velocity, the processivity exhibits a universal increase in response to external force. We use our results to analyze velocity and processivity data from single molecule experiments on the superfamily-4 ring helicase T7, and establish quantitatively that T7 is a weakly active helicase. We predict that compared to single-strand translocation, there is almost a two orders-of-magnitude increase in the back-stepping probability of T7 while unwinding double-stranded DNA. Our quantitative analysis of T7 suggests that the tendency of helicases to take frequent back-steps may be more common than previously anticipated, as was recently shown for the XPD helicase. Finally, our results suggest the intriguing possibility of a single underlying physical principle governing the experimentally observed increase in unwinding efficiencies of helicases in the presence of force, oligomerization or partner proteins like single strand binding proteins. The clear implication is that helicases may have evolved to maximize processivity rather than speed.

## INTRODUCTION

Helicases are enzymes that play an important role in almost every aspect of RNA and DNA metabolism [1–7]. This class of molecular motors, which has been grouped into six super families, utilize the free energy from ATP hydrolysis to translocate directionally over hundreds of bases on single-strand (ss) nucleic acids. In this respect, they are similar to other motors like kinesin and myosin, which also predominantly exhibit directional motion on microtubules and actin filaments, respectively [8, 9]. In addition, some helicases can also couple ss translocation with unwinding of double-stranded (ds) nucleic acids, thus playing a crucial role in cellular functions like replication, recombination and DNA repair. The “processive, zipper-like” ds unwinding activity was first demonstrated by Abdel-Monem and colleagues in *E. Coli* based DNA helicase in 1976 [10]. Interestingly however, most helicases cannot unwind double strands of nucleic acids by themselves, and require either oligomerization or assistance from other partner proteins to increase the efficiency of processive unwinding [11–13]. The fundamental principles governing this collaborative improvement of unwinding efficiency have not been fully elucidated.

A helicase that uses some of the energy of ATP hydrolysis to destabilize the ss-ds nucleic acid junction, is considered to be ‘active’. On the other hand, a helicase that utilizes the thermal fraying of the ds and opportunistically steps ahead when the site ahead is open, is referred to as ‘passive’ [2]. Considerable effort has been devoted to classifying various helicases into these two categories, as this general description gives an intuitive idea of the mechanism used by a particular helicase to unwind nucleic acids [14–24]. Earlier works used bulk kinetic assays or analysis of crystal structures to classify the activity of helicases [15–17], while more recent studies have primarily resorted to single-molecule experiments coupled with theoretical frameworks to establish the mode of unwinding of various helicases [20–23, 25–27].

Although ascertaining whether a helicase is active or passive is difficult and remains a controversial question for a number of helicases, we recently used this description in a quantitative framework to predict some surprisingly universal characteristics, which are independent of the dsDNA sequence [28]. By generalizing the original mathematical framework introduced by Betterton and Jülicher [29–31], we predicted that the processivity of active and passive helicases should always increase in response to external forces applied to the termini of dsDNA or dsRNA. Unlike the processivity, the velocity of unwinding, however, exhibits no such universal behavior, and exhibit a variety of responses to external force depending on whether the helicase is active or passive [28]. In our earlier work, we solved for the velocity and processivity of a helicase (with both the step-size and interaction range equal to one basepair) taking the sequence of DNA into account using numerically precise solutions. Here, we use a physically motivated approximation scheme to derive analytical expressions for both the velocity and processivity of a model that also accounts for a general step-size and interaction range. Similar models accounting for a non-zero step-size and interaction range have been used to study the velocity of unwinding using numerical simulations, without the benefit of easy to use analytical expressions [23, 32]. These expressions allow us to demonstrate the universality of the nature of the response of processivity to force, enabling us to draw far reaching conclusions on the evolution of helicases to optimize the number of base pairs that are unwound.

The analytic results derived here also allow us to analyze single-molecule experimental data on helicases more precisely than attempted before, by simultaneously analyzing the velocity and processivity of a particular helicase, instead of merely the velocity. The framework introduced by Betterton and Jülicher (and its variants), force or sequence dependence of the unwinding velocity had previously been fit to experimental data to determine the nature of T7, T4 and NS3 helicases [20–23]. However, Manosas *et al* [32] pointed out that the multi-parameter fit of the Betterton and Jülicher model to velocity data is not robust, with different parameter sets fitting the data equally well. More disturbingly, the various best-fit parameter sets suggest completely different unwinding mechanisms, and fail to provide any conclusive results for the fitting parameters. This is especially problematic when parameters like the step size and the potential back-stepping rate have not been characterized experimentally for a particular helicase, and need to be estimated from fits of the theory to data. As a consequence, the literature on helicases is rife with contradictory claims about the mechanism of action of a specific helicase. For instance, steady-state and pre steady-state kinetic assays [33] as well as studies of crystal structures [15] determined that NS3 helicase is passive, while a single molecule experiment coupled with a mathematical analysis suggested that NS3 is active [20]. Similarly, the T7 helicase was deemed to be an active helicase in two previous works [21, 23], while simple physical arguments suggested that T7 is passive [32]. Here, we show that our theory, which is used to analyze *simultaneously* velocity and processivity data can be used to robustly obtain the values of all the parameters associated with the unwinding activity of a helicase. Using data on the T7 helicase as a case study, we use our theory to explain both the force response of velocity and processivity, in the process quantitatively showing that T7 is very weakly active. Further, we recapitulate a number of independent experimental results on T7, thus confirming that the model is a good description of the helicase. Our theory also shows that an externally applied force, which *in vivo* could be generated by binding of single stranded partner proteins, results in an increase in processivity in all helicases, regardless of whether they are active or passive. This important finding prompts us to propose that helicases may have evolved to optimize processivity rather than speed.

## THEORY

### Model

The model, shown in Fig 1, is a generalization of the one introduced by Betteron and Jülicher [29, 31] that accounts for the finite processivity of the helicase, allows for an arbitrary step-size, and a general interaction range between the helicase and the double strand. Fig 1a shows the helicase (red filled circles) as it translocates on the ss nucleic acid strand (depicted as a bold black line). The position of the helicase on the nucleic acid track is denoted by *n*. The helicase can exhibit pure diffusion, and hence can step to the right or left with equal rate *k*^+^ = *k*^−^ = *k*. When the NTP hydrolyzes, the helicase moves forward at a rate *h* where *h* > *k*, thus resulting in a net forward rate *h* + *k* while the backward rate is *k*. If the mechanical step size of the helicase is *s*, then every time the motor steps forward or backward it does so by *s* nucleotides, resulting in an average velocity along the single strand given by *V_ss_* = *sh*. The helicase could also disengage from the nucleic acid track with a dissociation rate, *γ*.

**FIG. 1:**
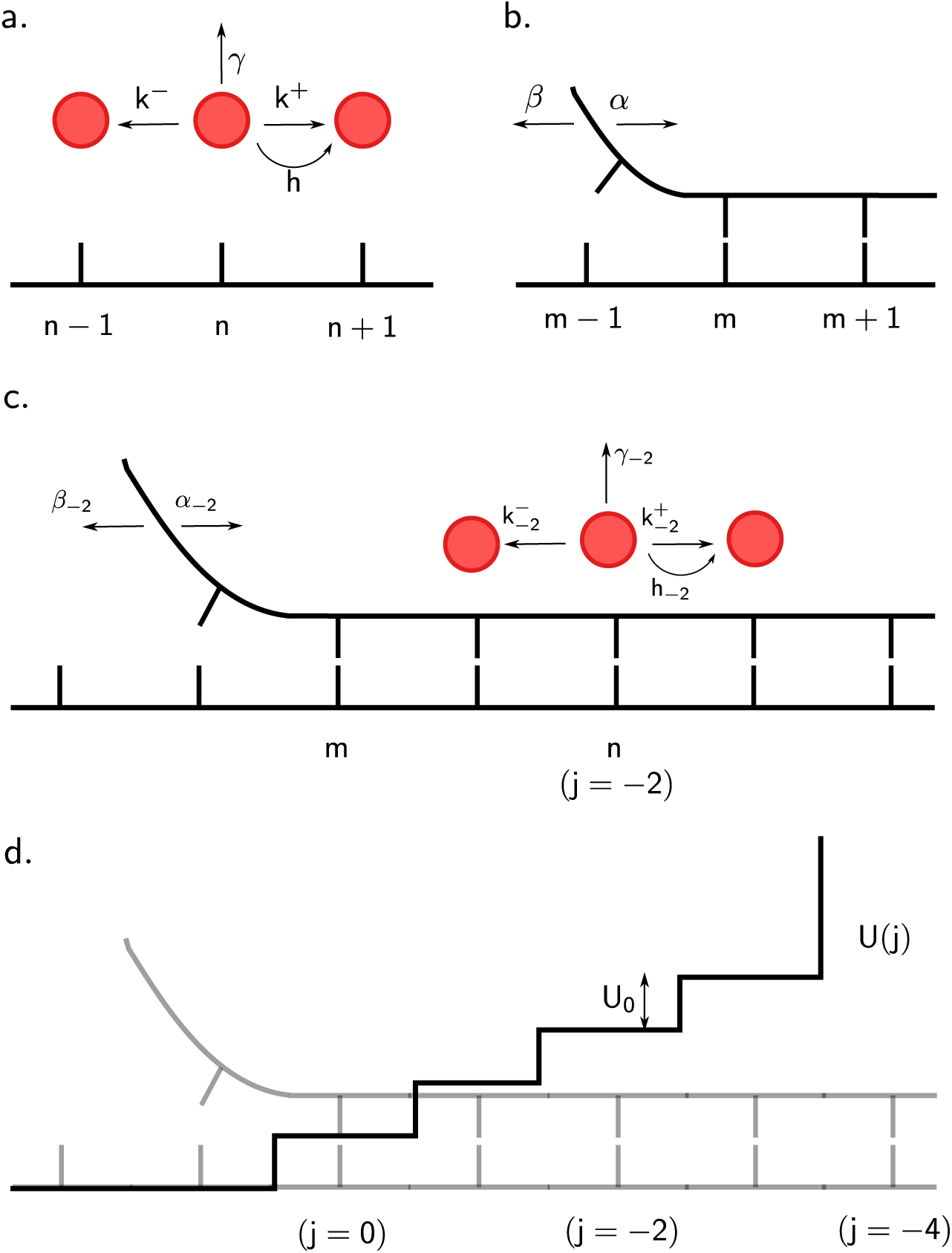
Model of nucleic acid unwinding by a helicase. (a) Single-strand stepping kinetics of the helicase. (b) Double-strand thermal ‘breathing’. (c) Modification of stepping kinetics of the helicase and breathing rates of the double strand. (d) The interaction potential that causes the modification of rates in (c).

Fig 1b represents the ss–ds junction at position *m*. The base pair at the junction can rupture at a rate *α* (increasing *m* to *m* + 1) while a new base pair can form (decreasing *m* to *m* − 1) at rate *β* such that *α/β* = exp(−Δ*G*), where Δ*G* is the stability of the particular ds base pair. All energies in this paper are in units of *k_B_T*.

Fig 1c shows how the rates change when the helicase and the junction approach each other. Modifications of the original rates occur due to the interaction potential *U*(*j*), a particular example of which is shown in Fig 1d. As the helicase and junction approach each other, they interact. As a result, the helicase has to perform extra work, which is *U*_0_ per base pair to step forward. This energy is provided from the hydrolysis of ATP. We follow the description in [30] and define *j* ≡ *m* − *n* to be the difference in the positions of the junction and the helicase. The rates are modified depending on *j*, and this is indicated by the value of *j* as a subscript. For example, *h*_−2_ denotes the modified forward stepping rate when *j* = −2 (Fig 1c). The motivation for allowing negative values of *j* and hence a potential as shown in Fig 1c, comes from the workings of helicases like PcrA and NS3 (see Fig 2b). These helicases seem to physically interact with downstream base pairs of the double strand [16, 20], possibly distorting and destabilizing a number of bases beyond the junction. Therefore, for a general scenario, we let the helicase interact with the ds over a range *r* of base pairs, after which a hard wall exists at *j* = −*r*. For ring helicases like T7, which encircle one strand of the DNA while excluding the other [34, 35], destabilization is not likely to happen by physical overlap with bases downstream of the junction (see Fig 2a), but might occur due to electrostatic interactions [36]. For such helicases, *j* would always be positive with a hard wall at *j* = 0. Note that the exact position of the potential does not matter: a shifted potential with the first step at *j* = 4 and hard wall at *j* = 0 (Fig 2a) would give identical results for the velocity and processivity as the potential shown in Fig 2b. Hence this model for the interaction range of the helicase is general. The short range potential is chosen to have a constant step height *U*_0_ and is defined as follows:

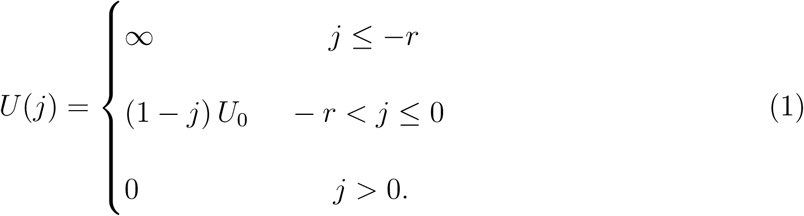

**FIG. 2:**
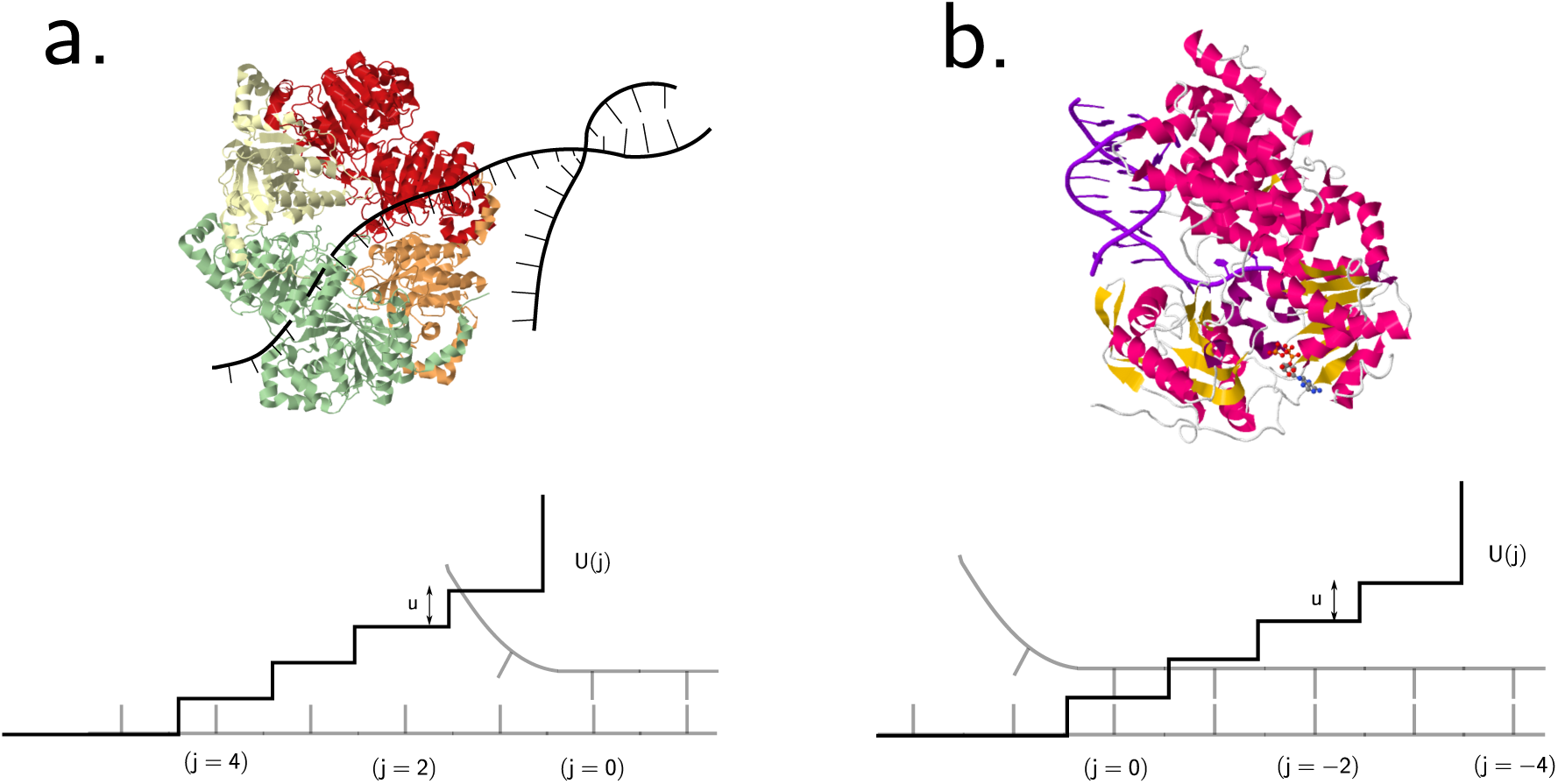
Different scenarios for the interaction potential between the helicase and double-strand. (a) Crystal structure of T7 helicase (Protein Data Bank (PDB) code 1E0J) and cartoon of ds DNA. The helicase encircles one strand and excludes the other, suggesting that the interaction with the junction must happen from a distance, presumably due to electrostatic forces. (b) Crystal structure of the PcrA helicase bound to ds DNA (PDB code 3PJR), showing the overlap of the helicase domains with bases beyond the junction. Below both the figures are shown the corresponding interaction potentials *U*(*j*). Identical expressions for the unwinding velocity and run-length result for both the potentials.

The nucleic acid breathing rates are modified because of this interaction as follows:

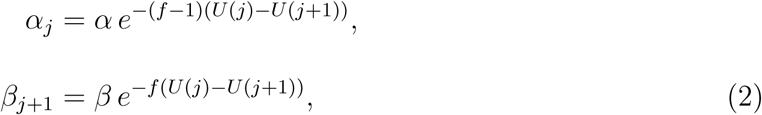
such that 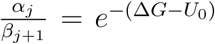 for all values of *j*. In Eq. 2, *f* is a number between 0 and 1 representing the fractional position of the free energy barrier between base-pair open and base-pair closed states, from the open state. Nucleic acid opening takes the system from state *j* to state *j* + 1 while closing takes the system from *j* +1 to *j*. Hence the exponents in Eq. (2) involve the term *U*(*j*) −*U*(*j* + 1). The rates of the helicase also get modified because of the interaction, and change in the following way:

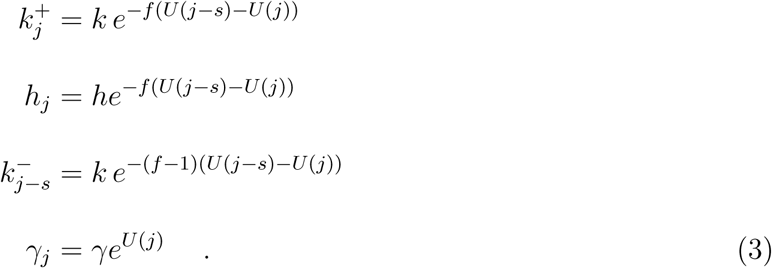

With a step size *s*, the helicase cannot move to the right if *j* ≤ *s* − *r*, and hence 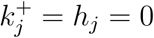 for all *j* ≤ *s* − *r*. Eq. (3) shows that as long as the helicase and the ss–ds junction are separated by a distance *j* > *s*, the forward rates are independent of 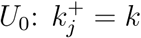 and *h_j_* = *h*. For the backward rate, 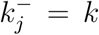 as long as *j* > 0. Notice that the exponents in Eq. (3) contain the term *U*(*j* − *s*) − *U*(*j*) since the helicase jumps *s* nucleotides every time it steps forward or backward.

### Force effects

To model the effect of a constant external force *F* applied directly to the ds junction, the ds opening and closing rates change from *α_j_*, *β_j_* to 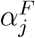, 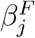 such that:
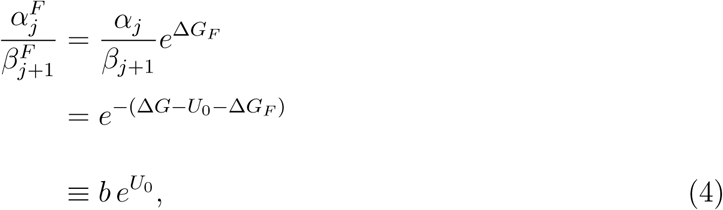
where we defined 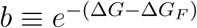, and Δ*G_F_* is the destabilizing free energy of the base pair at the junction, due to the constant external force *F*. Assuming a Freely-Jointed-Chain model for the single strand DNA segments, the expression for Δ*G_F_* is given by [37]:
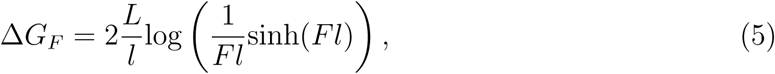
where *L* is the contour length per base and *l* is the Kuhn length. In writing Eq. (4), we assume that all of the external force *F* is transmitted through the single strands and destabilizes the base pair at the junction. For ring-shaped helicases like T7, which encircle one strand while excluding the other [34, 35], this model is a very good description. However, this may not be an accurate description of other helicases like the NS3, where the domains surround both the strands of the nucleic acid, and the junction may be protected to some extent from external forces [5]. For such helicases, a more careful analysis is needed, and is left for future work.

With the model thus defined, we need to solve for the velocity and the processivity of the helicase as it unwinds the ds nucleic acid. The velocity is defined as the average number of bases per unit time that the helicase moves to the right, in a binding event. For the processivity, multiple definitions have been proposed [31]. The mean binding time 〈*τ*〉, the translocation processivity 〈*δn*〉 which gives the average distance moved by the helicase in a binding event, and the unwinding processivity 〈*δm*〉 which gives the distance moved by the ss–ds junction during a single binding event of the helicase–are used as measures of helicase processivity. As shown numerically in [31], the latter two definitions of processivity are almost identical when the helicase attaches close to the ss–ds junction. Since this is the physically relevant situation, as we argued previously [28], we will not differentiate between 〈*δn*〉 and 〈*δm*〉 in this work, and refer to both as the run-length or processivity.

## UNWINDING VELOCITY AND RUN-LENGTH: SOLUTION OF THE MODEL

To solve for the velocity and run-length of a finitely processive helicase, we first note that *α* and *β* are very large [38–40]–larger by orders of magnitude compared to any rate describing the kinetics of the helicase. Therefore, even before the helicase takes a single step (backward, forward or detach), the ss–ds junction would have opened and closed multiple times. As a result, the probability *P_j_* of observing the helicase and junction at a separation *j* would have reached a steady state distribution, long before the helicase moves. Since there is a hard wall at *j* = −*r*, there is no probability current between *j* and *j* + 1 in this steady state, for any value of *j*. This, along with the normalization condition, 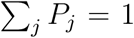, allows us to solve for *P_j_*:

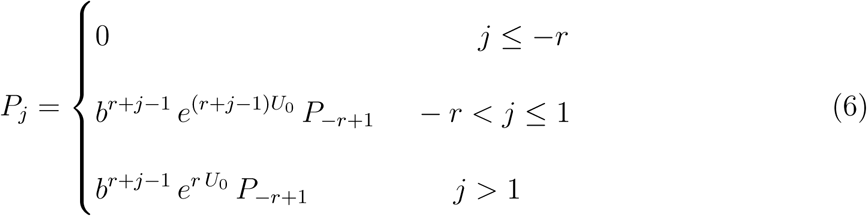
where,

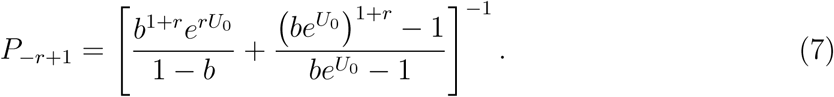
where *b* as defined before, is given by 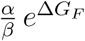. The two-body problem of the helicase and junction can now be recast in terms of a one-body problem involving only the helicase, moving with renormalized rates that we denote with a tilde 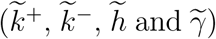; 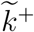 is given by 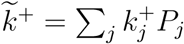, and similar expressions describe all the other renormalized rates. Performing the sums, the final lengthy expressions for 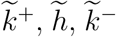 and 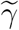 are given in the Appendix. Notice that although *k*_+_ = *k*_−_ = *k*, the helicase–junction interaction causes the renormalized rates to become different, hence 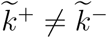.

The unwinding velocity is given by:

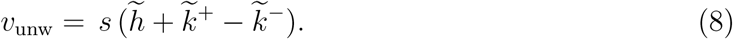

For *s* = 1 and *r* = 1, *v*_unw_ reduces to,

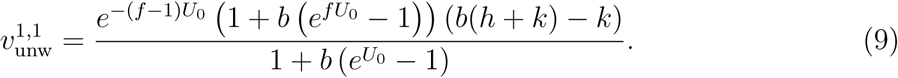

The expression for one-step active unwinding derived by Betterton and Jülicher (Eq.(27) in [30]) reduces to our expression in Eq. (9), when all the rates associated with the helicase are neglected compared to *α* and *β*.

The mean attachment time of the helicase 〈*τ*〉 is given by the inverse of the renormalized detachment rate:

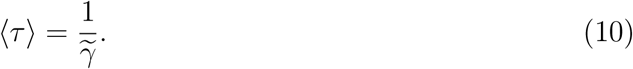

The processivity is given by 〈*δm*〉 = *v*_unw_〈*τ*〉. Using Eq. (8) and Eq. (10), we obtain,

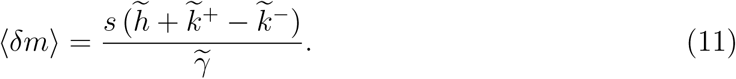

For *s* = 1 and *r* = 1, 〈*δm*〉 reduces to the following expression:

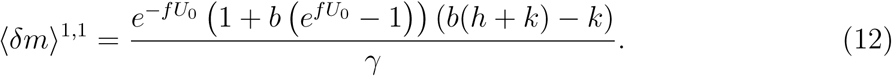

Eq. (8), Eq. (10) and Eq. (11) along with Eq. (16) are the important results in this paper. Note that by defining *k*^−^ ≡ *k* and *k*^+^ ≡ *k* + *h* in Eq. (9) and Eq. (12), we obtain the same model used in our previous work [28].

## UNIVERSAL FORCE RESPONSE OF THE UNWINDING PROCESSIVITY

In our earlier work [28], we showed numerically that the unwinding velocity and processivity show contrasting responses to external force. We analyzed a model with step size of one basepair and an interaction potential with a one basepair range. Below, we first revisit the earlier model to highlight the basic results and show quantitatively the differences between the responses of velocity and processivity to force. We then point out the effects of increasing the step-size and the interaction range of the helicase.

For *s* = 1 and *r* = 1, the expressions for velocity and processivity are given by Eq. (9) and Eq. (12) respectively. As in our previous work, we choose Δ*G_F_* = *F*Δ*X* for illustrative purposes. Choosing this simple form instead of the more accurate model based on the Freely Jointed Chain (Eq. (5)), does not qualitatively change any of the results [28]. We will look at the limit *k* = 0, to simplify all the analytic expressions. For the more general case of non-zero *k*, as long as *k* ≪ *h*, all the results are valid. Helicases are believed to satisfy this criterion [32], and is supported by our fitted values (discussed below) from data on the T-7 DNA helicase.

Setting Δ*G_F_* = *F*Δ*X*, and differentiating Eq. (9) with respect to *F*, we obtain the following expressions for passive (*U*_0_ = 0) and optimally active (*f* = 0*,U*_0_ = Δ*G*) helicases. Note that optimally active helicases are ones where *f* → 0 and *U*_0_ ≥ Δ*G*.

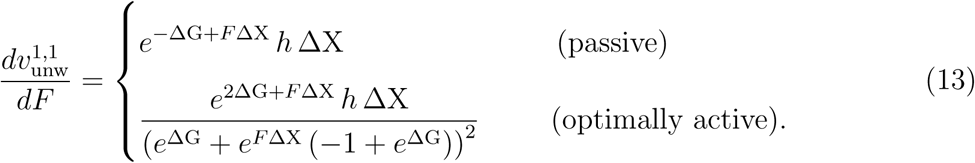

The equation above shows that for a passive helicase, the slope of the velocity-force curve is not only always positive, it increase exponentially with *F* (orange curve in Fig. 3a). On the other hand, for an optimally active helicase, the expression for the slope has the term *e^F^*^Δ^*^X^* both in the numerator as well as the denominator–with a higher power of *F* in the denominator, implying that the increase in velocity with force will be much less rapid than an exponential. As can be seen from Fig. 3a, the curves (green, blue and red) have the opposite curvature compared to the passive helicase (orange curve). In contrast, the force-dependent behavior of the processivity shows a universal increase, as can be seen from the following equation:

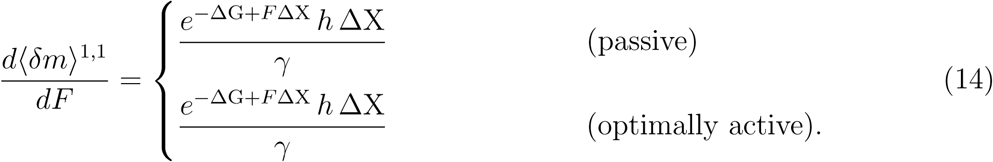

**FIG. 3:**
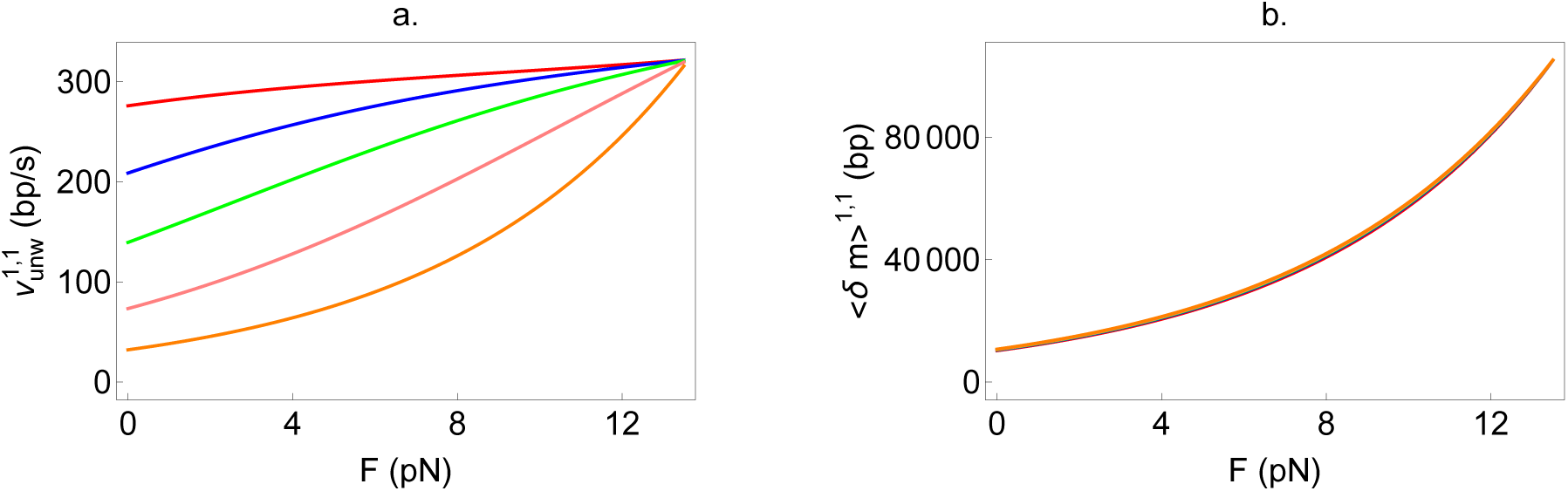
Response of (a) velocity and (b) processivity to external force, for a helicase with a 1 bp step-size and a 1 bp interaction range. The orange curve corresponds to a passive helicase while the red, blue and green curves correspond to optimally active helicases. The velocity curves are plots of Eq. (9) while the processivity curves are plots of Eq. (12).

Eq. (14) shows that the slopes are identical for both passive and optimally active helicases, and increase exponentially with force. This is very clearly illustrated in Fig. 3b, where all the curves almost superpose.

For a general step-size and interaction range of the helicase, the full expressions for unwinding velocity (Eq. (8)) and processivity (Eq. (11)) are complicated, and hence we only show a few representative plots in Figs. 4 and 5, for a variety of step-sizes and interaction ranges. Fig. 4 shows that increasing the range of interaction (keeping the step-size fixed) affects the velocity and processivity in different ways. At a given force, while the velocity of unwinding increases when the range is increased (Fig. 4a,b), the processivity decreases (Fig. 4c,d). However, the universal behavior obtained in our previous work [28] remains valid: the unwinding velocity can increase or remain almost constant with force, depending on whether the helicase is active or passive. The processivity on the other hand, always increases with external force. Fig. 5 shows the effect of increasing the step-size while keeping the interaction range fixed, for optimally active (Fig. 5a,c) as well as very weakly active (Fig. 5b,d) helicases. At a given value of force, both the velocity and processivity decreases with increase in the step size. For an optimally active helicase, the decrease in velocity is rapid, becoming strongly negative at low forces (Fig. 5a). The reason for this phenomenon is that the back stepping rate becomes larger as *U*_0_ is larger (more active helicase). The helicase-junction interaction leads to the helicase imparting a force on the junction (= *U*_0_ divided by the base-pair distance). This in turn causes an opposite force from the junction on the helicase. This opposing force is larger for larger *U*_0_, thereby leading to larger effective back-stepping rates as *U*_0_ increases. For larger step-sizes (3 bp for the green line in Fig. 5a), the helicase needs more bases open downstream simultaneously and hence is less likely to step forward. Coupled with a larger back-stepping rate, the net effect is negative unwinding velocities of the helicase.

**FIG. 4:**
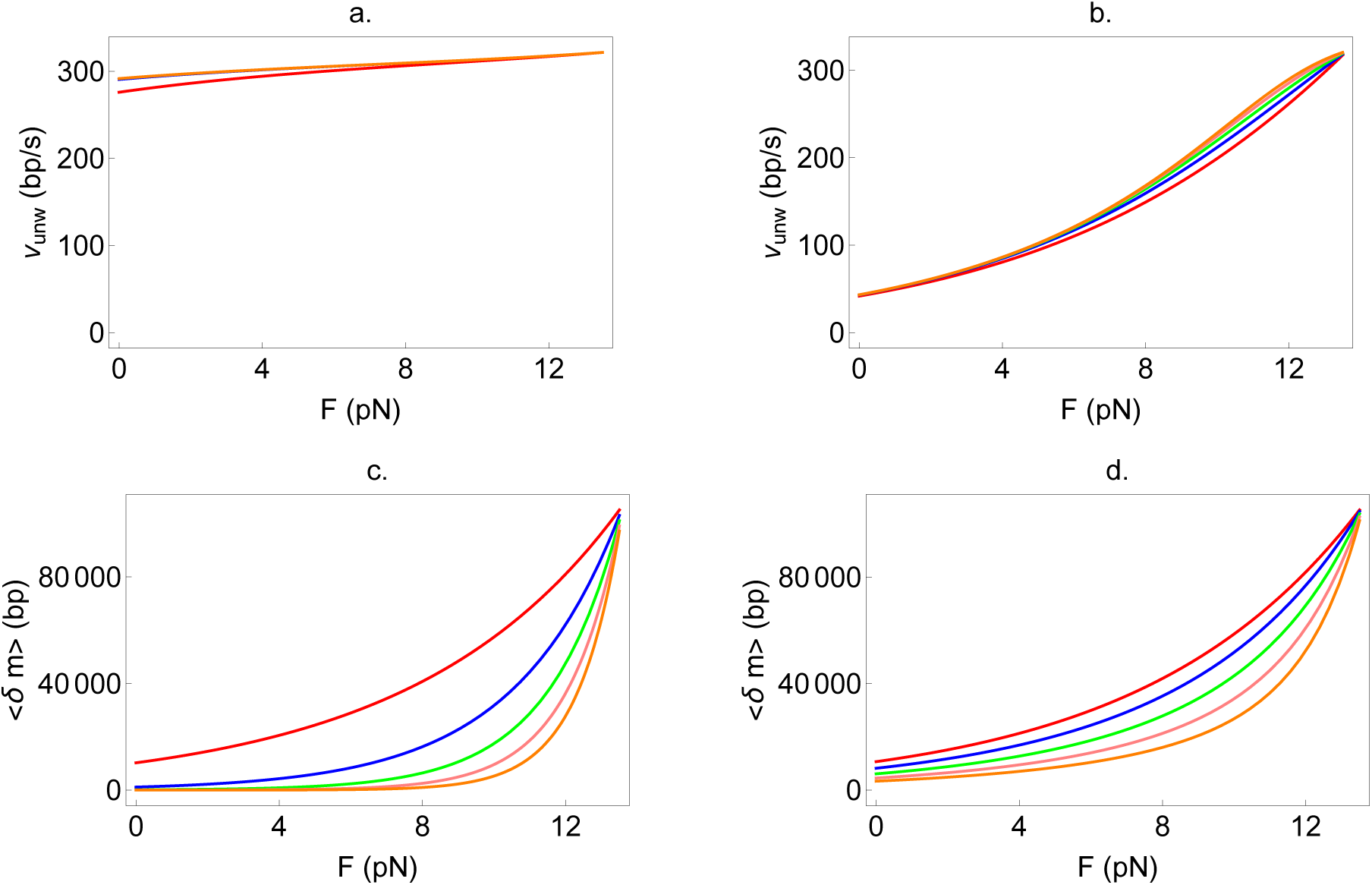
Effect of increasing the interaction range of the helicase on the unwinding velocity and processivity, while keeping the step-size fixed. The step size in all panels is fixed at 1 bp and *h* = 322*s*^−1^, thus *V_ss_* = 322 bp/s. The interaction ranges are 1 bp (red), 2 bp (blue), 3 bp (green), 4 bp (pink), 5 bp (orange). (a) and (c): Optimally active helicase with *U*_0_ = 5 *k_B_T* and *f* = 0.01. (b) and (d): Very weakly active helicase with *U*_0_ = 0.3 *k_B_T* and *f* = 0.01.

**FIG. 5:**
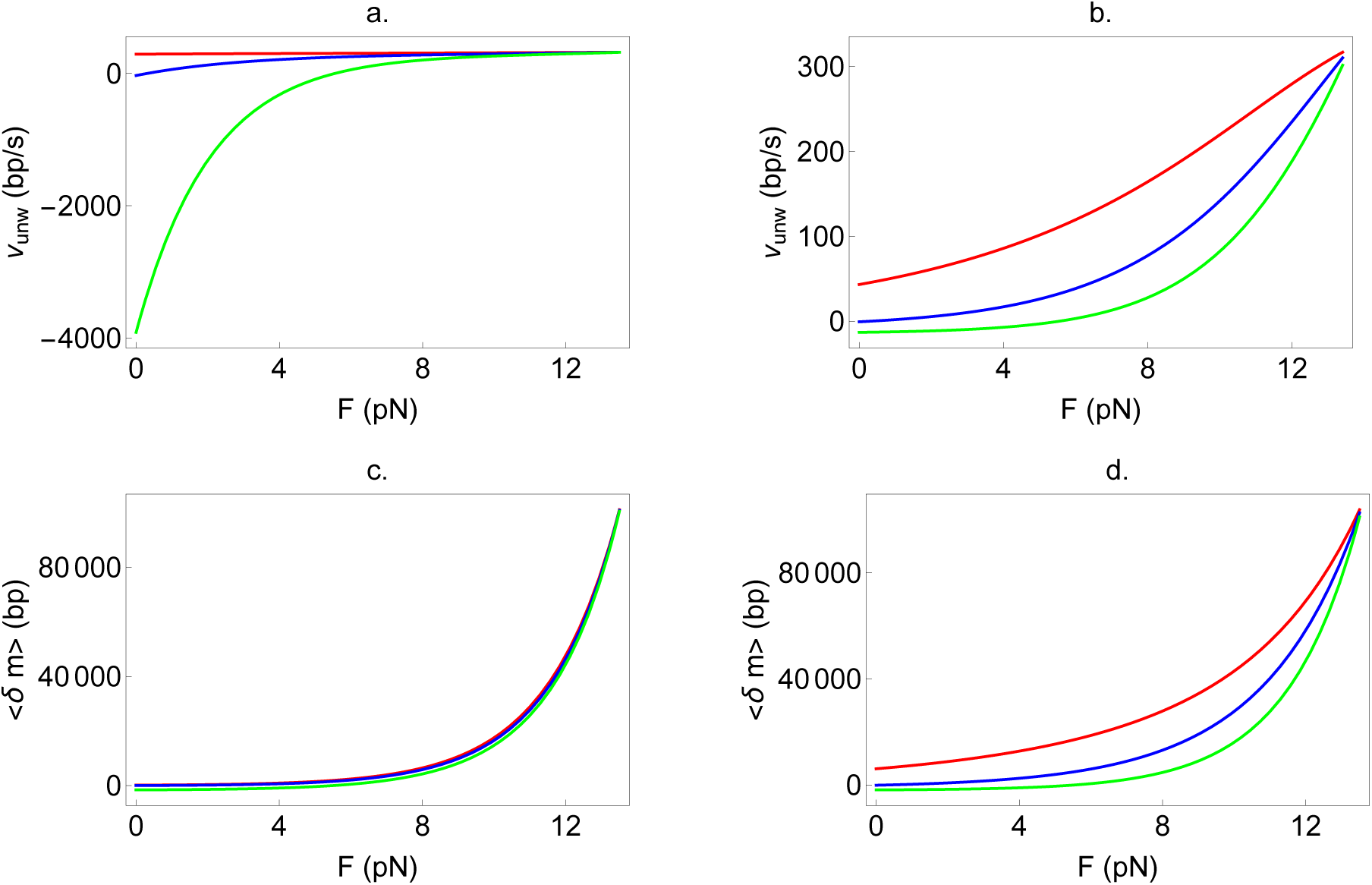
Effect of increasing the step-size of the helicase on the unwinding velocity and processivity, while keeping the interaction range fixed. The interaction range in all panels is fixed at 3 bp and 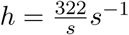, thus *V_ss_* = 322 bp/s. The step-sizes are 1 bp (red), 2 bp (blue), 3 bp (green). (a) and (c): Optimally active helicase with *U*_0_ = 5 *k_B_T* and *f* = 0.01. (b) and (d): Very weakly active helicase with *U*_0_ = 0.3 *k_B_T* and *f* = 0.01.

To summarize, our analysis in this section shows that irrespective of the step-size, interaction range, active or passive nature of the helicase, the unwinding processivity should always increase with force. The unwinding velocity however does not exhibit the same universal increase with external force.

## UNWINDING MECHANISM OF THE T7 HELICASE

### Simultaneous fitting of DNA unwinding velocity and run-length data

Since fitting only the expression for the unwinding velocity to experimental data proves to be insufficient for estimating physically reasonable parameters, we reasoned that fitting the theory to the available data to the two observables simultaneously should significantly limit the parameter space and allow for better extraction of the important parameters of the system. Using the velocity given in Eq. (8) and the processivity given in Eq. (11), we use our theory to analyze data from the T7 helicase.

To apply our theory to analyze the data, we first observe that the velocity as a function of force has the following parameters: *U*_0_*, k, f, s, r* and *h*. It follows from Eq. (11) that the processivity has one extra parameter, the dissociation rate *γ*. However, *γ* is a quantity that is measured in bulk experiments [41, 42], while the relation *V_ss_* = *sh* allows us to use the experimentally determined value of *V_ss_*, to reduce another free parameter. Thus, the number of free parameters left to be determined from fitting to experimental data is five—*U*_0_*, k, f, s* and *r*. To test our method, we fitted velocity and processivity data from single molecule experiments on the T7 helicase (Fig 6b and Fig S6 respectively, of [23]). Kim *et al* [41] reported *γ* = 0.002*s*^−1^ at 18°C. Since the single molecule experiment was performed at 25°C, we used the rough estimate that around room temperature, a number of chemical rates increase by about a factor of 2–3 for every 10°C increase [43], to estimate *γ* at 25°C. We therefore used *γ* = 0.003, 0.004 and 0.005*s*^−1^. We used *h* = 322/*s* (*s* is the step size) since the ss velocity was measured to be 322 bp/s [23], which seems not inconsistent with the bulk result of 132 bp/s at 18°C [41]. We took Δ*G* = 2.25 since the DNA sequence had 48% GC content (supplementary information of [22]). Both the bulk and single molecule experiments were performed at 2 mM dTTP concentration.

**FIG. 6:**
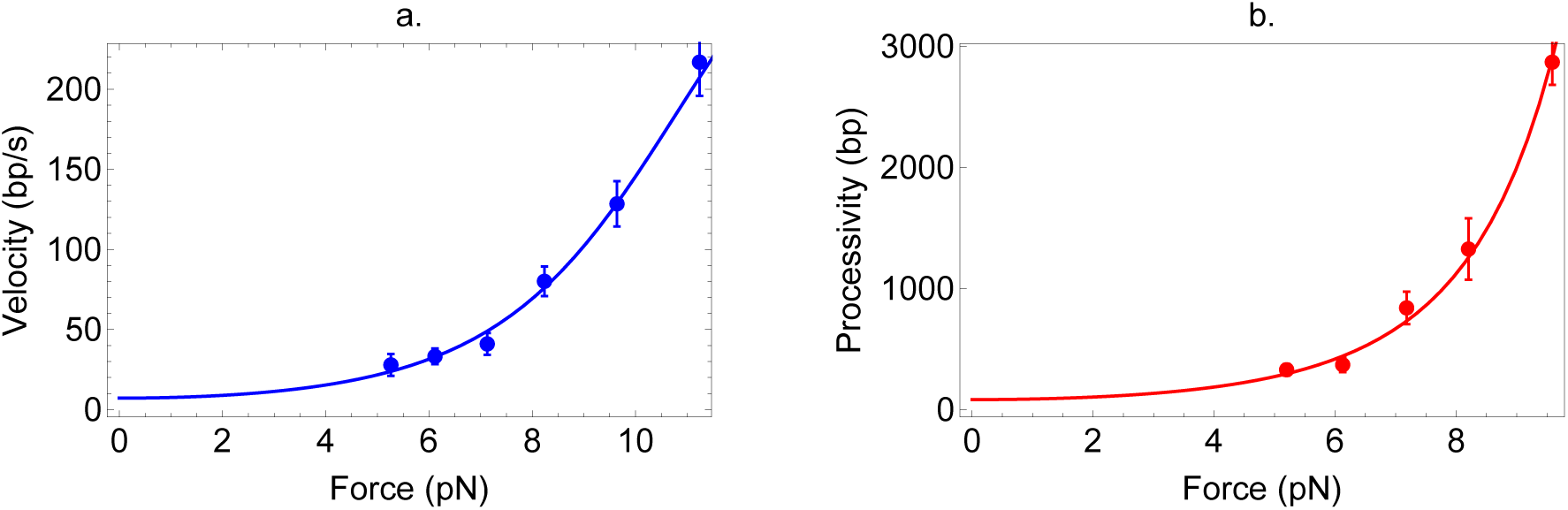
Simultaneous fitting of velocity and run-length data. The blue and red circles are experimental data for the T7 helicase from [23]. The blue line in (a) is Eq. (8) fitted to velocity data while the red line in (b) is Eq. (11) fitted to processivity data. The best fit parameters are given in Table II.

For the force dependent destabilization of the double strand given in Eq. (5), the parameters *L* and *l* need to be chosen carefully, to reproduce the critical force *F_c_* observed in the experiment. *F_c_* (the force where Δ*G* = Δ*G_F_*) was observed to be around 13.6–13.7 pN for the ds DNA sequence we have analyzed in this work (Fig 6b of [23]). The usual values chosen for *L* and *l* are 0.56 and 1.5 nm respectively [37]. However, *F_c_* for this choice of parameters (and Δ*G* = 2.25) is about 15 pN, so we chose *L* = 0.63 nm and *l* = 1.5 nm to reproduce the observed critical force. To check the robustness of our results, we also tried a different parametrization *L* = 0.56 and *l* = 1.95 nm, which results in *F_c_* = 13.6 pN. Both these parametrizations produce nearly identical results, hence we show results with only the first one (Table I and II).

**TABLE I:** Fitting to only velocity data. Shown are just a few fits, obtained by varying *s* and *r*, from all those that are similar. The definition of ‘similar’ is that the worst fit should be 0.5 times as likely as the best fit, according to AICc (see text for details).

| Step Size $s$ (bp) | Interaction Range $r$ (bp) | $\chi^2$ |
| --- | --- | --- |
| 1 | 20 | 3.04 |
| 2 | 10 | 1.65 |
| 2 | 20 | 1.72 |
| 2 | 50 | 1.74 |
| 3 | 3 | 2.3 |
| 3 | 4 | 2.8 |

**TABLE II:**
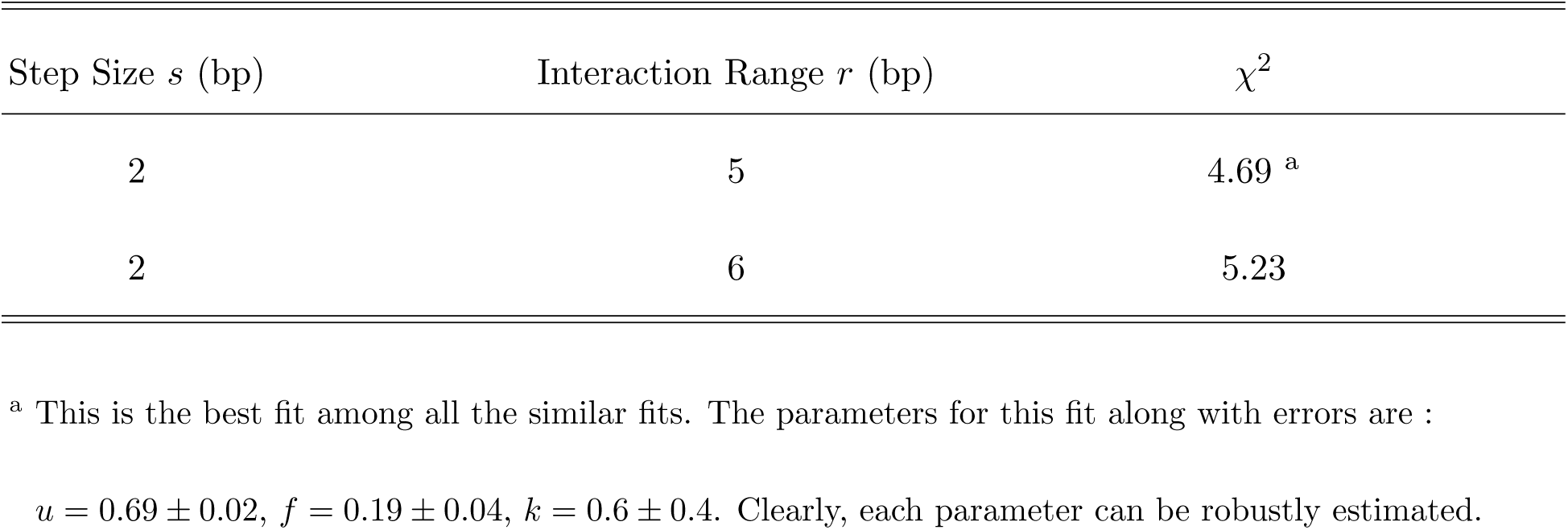
Fitting simultaneously to velocity and run-length data. Compared to Table I, the parameter space with similar fits to the data has been drastically reduced. The definition of ‘similar’ is exactly the same as that used in Table I.

### Quality of fits

The results of simultaneous fitting are shown in Fig 6. Tables I and II show the quality of fits (*χ*^2^) for a variety of parameter sets with ‘similar’ fits. To quantitatively define ‘similarity’ of fits, we used the Akaike Information Criterion (AICc) [44] defined as:

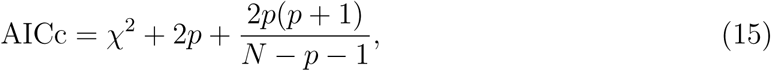
where *N* is the number of data points and *p* the number of free parameters. The usefulness of this criterion is that the quality of the two sets of fits can be quantitatively compared: if two model fits have AICc values of *a*_1_ and *a*_2_ respectively, with *a*_1_ < *a*_2_, then model 2 has a likelihood exp((*a*_1_ − *a*_2_)/2) of being the true interpretation of the data, relative to model 1. Using this interpretation, we show in Tables I and II all fits that are at least 0.5 times as likely as the best fit among that set. Table I shows the result of fitting to only velocity data—multiple parameter regions can fit the velocity data with similar quality of fits. Table II shows results of our simultaneous fitting procedure. Clearly, Table II shows that the simultaneous procedure allows a much narrower range of parameters to produce similar fits. In addition, the errors on the parameters are small, allowing all five parameters to be extracted with great robustness. The best fit parameters are *U*_0_ = 0.69 *k*_B_*T*, *f* = 0.19, *k* = 0.6*s*^−1^, *s* = 2 bp and *r* = 5 bp.

### Comparison with experiments on T7 unwinding of DNA under zero force conditions

An earlier bulk experiment [45], and a more recent FRET-based single molecule experiment [46] on T7 DNA, were carried out under conditions of zero external force. Using an ‘all-or-none’ assay at 18°C, five DNA sequences (average GC content of 37%) were unwound with T7 in [45], resulting in an average unwinding velocity of 15 bp/s. Approximately consistent with these results, the unwinding velocity at 23°C of T7 on a 35% GC sequence was found to be 8 bp/s [46]. Fig 6a shows our model prediction for the unwinding velocity at zero force: *v_unw_* = 7.1 bp/s. Keeping all the parameters fixed at the values shown in Table II, but reducing Δ*G* to 1.9 to correspond to a DNA sequence comprising roughly 37% GC basepairs, our model predicts an unwinding velocity of 18 bp/s at zero force. Taking into account that our analysis is based on an experiment performed at a slightly higher temperature (25°C) compared to either of these two zero-force experiments, our results are consistent with the two previous works.

### Predictions for sequence dependence of detachment and back-stepping rates of T7 while unwinding dsDNA at zero-force

The sequence dependence of the detachment rate of a helicase is an aspect that can be directly measured in experiments [20, 47], and can be an indicator of whether the helicase is active or passive. By fixing the parameters in our model to the best-fit values of Table II, and changing only Δ*G*, we can predict how the detachment and back-stepping rate of T7 will depend on the sequence composition, while unwinding DNA. The results of this analysis is shown in Fig 7. As is evident, neither the detachment rate, nor the back-stepping rate are very sensitive to Δ*G*, because the helicase is only very weakly active. Interestingly, this is akin to the observations made in a previous experiment on the DnaB helicase [47], where it was shown that for sequences with 50–100% GC composition, the detachment rate was almost constant. Since both DnaB and T7 are very similar in structure and sequence, belonging to the superfamily-4 group of helicases, our results suggest that the ring helicases of the superfamily-4 group might all use a similar weakly active mechanism for unwinding DNA.

**FIG. 7:**
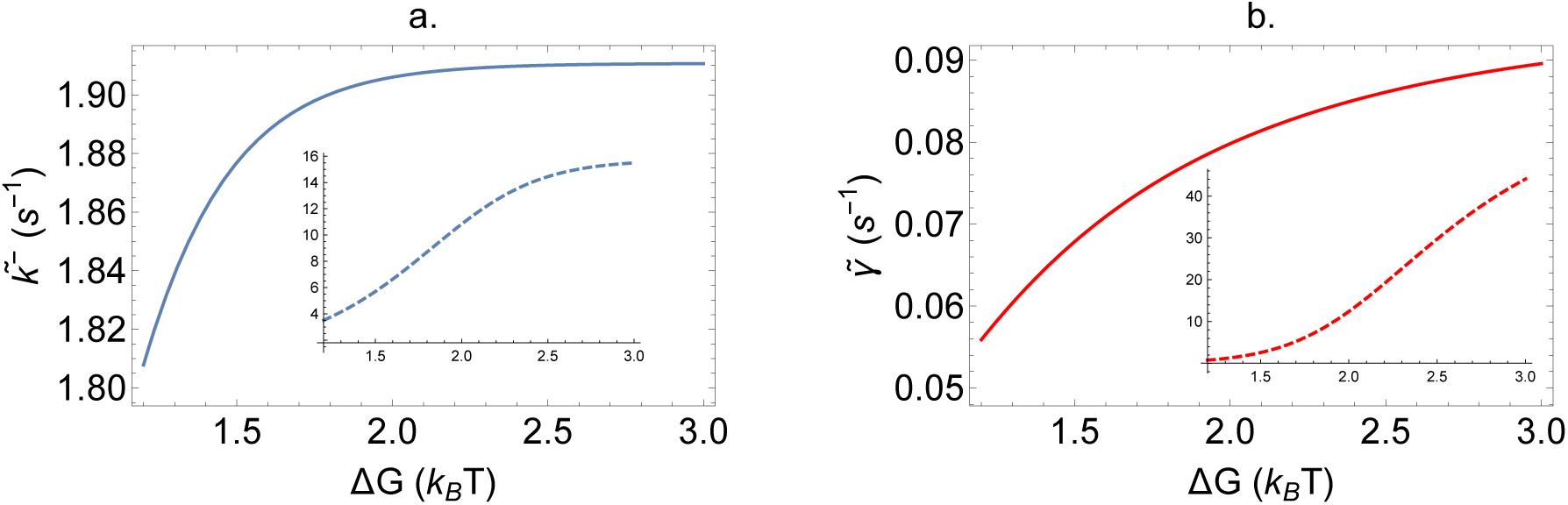
Predictions for sequence dependence of T7 detachment and back-stepping rates while unwinding ds DNA. (a) The back stepping rate while unwinding 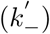 hardly changes as a function of the sequence stability, as a result of the helicase being only weakly active. (b) Similarly, the detachment rate while unwinding 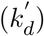 changes by only a factor of 1.5 with change in Δ*G*. The insets in both figures show the hypothetical situation of a highly active T7, with *U*_0_ = 2.0 *k_B_T*. Both the back-stepping and detachment rate show much more sensitivity to Δ*G* under highly active circumstances.

## DISCUSSION

### T7 is weakly active

From the best fit parameters in Table II, we infer thatthe T7 helicase is a weakly active helicase, destabilizing the ds junction by about 0.69 *k*_B_*T* per base. The value of *U*_0_ obtained here, is different from the (1 − 2)*k*_B_*T* estimate reported earlier in [21, 23]. Both these works analysed only velocity data – the former looked at sequence dependence while the latter examined the force dependence of velocity. It was originally pointed out [32] and verified by us specifically for T7 in this work (Table I), that analyzing only the velocity data using the multi parameter Betterton and Jülicher model is not sufficient for robust parameter estimates. There are other differences as well between our model and the ones used previously [21, 23]. The parameter *f* was fixed to 0.05 in both those earlier studies, whereas we allow it to vary, given that *f* is a physical quantity which could take on any value between 0 and 1. We also have the extra parameter *k*, which gives the rate of pure diffusion. The presence of this parameter allows for back-steps, which was neglected in the previous studies. It is important to include this parameter, especially in light of recent experiments that directly observed back-stepping [24, 46].

### Step-size of T7

Our prediction of a step-size of 2 bp (2 bases advanced for each ATP hydrolyzed) is in agreement with certain previous experimental results. Using a pre steady-state analysis, it was found that one ATP molecule is consumed for every 2 − 3 basepairs translocated by T7 on a ss DNA [41]. A crystal structure of the DnaB helicase bound to ssDNA, showed that the step-size of DnaB is 2 bp [48]. DnaB and T7 are both members of the superfamily-4 group of helicases, with very similar sequence and structure in the C-terminal domains [19, 49]. These results suggest that the step size of T7 while unwinding ds DNA may also be 2–3 bp, under the assumption that ss translocation and ds unwinding occur with the same step-size. A recent smFRET-based unwinding assay using T7 observed stochastic pauses after every 2–3 bp of G-C rich DNA unwound [46]. However, the waiting times of these pauses were gamma distributed rather than exponentially distributed, suggesting the presence of hidden steps within those pauses. Though the results do not prove a direct association of these hidden steps with ATP consumption, it would not be surprising if the helicase has a distribution of step-sizes with shorter steps of 1 bp while unwinding G-C bases. Since our model does not distinguish between the step size during translocation, unwinding or for different sequences, it is likely that our result of 2 bp per ATP consumed is a reflection of the average step-size over the entire ds sequence of the DNA being unwound. Further experiments, including determination of crystal structures, would be able to shed more light on this interesting conundrum.

### Interaction range of T7

The interaction range of ~ 5 bases that we obtain from our fits, is physically reasonable given the structure of the T7 ring helicase and its mode of binding to dsDNA. While the DNA strand excluded from the T7 ring is negatively charged, the C-terminal face of T7 is also negatively charged [36]. Replacement of the charged residues on the C-terminal by uncharged ones leads to a reduction in efficiency of complementary strand displacement [50]. These results strongly suggest that the moderately weak (0.69 *k_B_T* per base) interactions between the helicase and DNA that we predict, are electrostatic in nature. Given that the Debye-Hückel screening length is ~ 1 nm in physiological salt concentrations [51], the range of electrostatic interaction of the helicase should be a few nanometers. Our prediction of 5 bases (~ 1.7 nm) therefore is physically reasonable. Notice that fitting to only velocity data would predict an interaction range greater than 20 bp, which would be unphysically large.

### Back-stepping rate of T7 while unwinding or translocating

The back-stepping rate of helicases is usually very difficult to measure directly, due to lack of sufficient resolution in single-molecule experiments. Hence, the back-stepping rate is usually assumed to be negligible compared to the forward stepping rate or neglected completely [32]. Using the parameters extracted from our fits to T7 data (given in Table II), we now show that though this assumption is valid when the helicase translocates on ssDNA, the back-stepping probability is orders of magnitude larger while unwinding dsDNA. Note that in this discussion and throughout this paper, we set the ATP concentration at 1mM, which is roughly the physiological concentration of ATP. Experiments are usually performed at ATP concentrations of 1mM or higher. If the ATP concentration is made very low in an experiment, naturally the back-stepping probability will be substantially larger [24] until it approaches 50% at 0 ATP, implying no directionality associated with the motion of the helicase.

The ratio of forward to backward stepping rates while translocating is (*h* + *k*)/*k*, hence using *k* = 0.6 and *h* = *V_ss_*/*s* = 161, this ratio turns out to be 269. This result is interesting, and suggests that T7 back-steps approximately as frequently as some of the other processive molecular motors like Kinesin, which also has similar values for this ratio in the absence of external force [52]. The back-stepping probability at every step is given by *k*/(*k* +(*h* + *k*)), which is a mere 0.3% for the T7 parameters given in Table II. However, when the helicase unwinds dsDNA, the rates change, and the modified back-stepping probability is given by 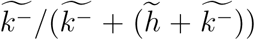. For the same parameter set, this works out to be 26%, almost two orders of magnitude larger than the back-stepping probability while translocating. Our prediction regarding this enhanced back-stepping probability while unwinding, is similar to the observations in a recent experiment on the XPD helicase, where it was shown that at 1mM ATP the back-stepping probability is about 10% [24]. Our analysis therefore suggests that XPD, which belongs to superfamily 2, may not be unique in this respect – the superfamily 4 helicase T7 also seems to back-step with relatively large probability while unwinding dsDNA. Further experiments on helicases belonging to the same super family are needed to probe the extent of back stepping.

### Universal nature of force response of helicase processivity, oligomerization and partner proteins

Helicases often cannot unwind double strand nucleic acids by themselves, but require partner proteins like single strand binding proteins or oligomerization to increase the efficiency [11–13]. We use efficiency to mean that the helicase is highly processive. While the mechanistic reasons for the increase in efficiency is likely to vary for individual helicases, our work suggests the possibility of a single physical principle underlying this effect. Our theory shows that the increase in processivity in response to external force holds for all helicases. In particular, unlike the unwinding velocity, the processivity increases with external force irrespective of the active or passive nature of the helicase. Since the external force destabilizes the double strand and decreases the free energy of the base pairs at the junction, we predict that any perturbation that results in reduction of base-pair stability should result in universal effects, similar to the force response of helicases. There is strong evidence that partner proteins, like single strand binding proteins (SSB), melt nucleic acids by reducing the stability of base pairs (see a more detailed discussion in [28]). Similar effects may be achieved by oligomerization of helicase monomers. A recent study on the UvrD helicase showed that dimerization leads to a closed conformation of the 2B subdomain of the leading UvrD monomer[13], a conformation that contacts the junction duplex and presumably destabilizes it [53, 54]. This implies the possibility that a universal underlying principle governs the response of helicases to force, partner proteins as well as oligomerization. An immediate prediction of this theory is that a weakly active helicase like T7 should exhibit increased velocity and processivity in the presence of partner proteins. This phenomenon has indeed been observed previously, but not explained [55, 56]. On the other hand, partner proteins should increase only the processivity of an active helicase like NS3. This prediction is borne out as well in experiments [57]. It is rare to find universal behavior, especially on the *nm* scale representing motors including helicases. That this seems to be the case for helicases, is remarkable. The universal increase of processivity with force, regardless of the nature of the helicase, also suggests that this class of motors may have evolved to optimize processivity rather than speed.

## CONCLUSIONS

Whether a particular helicase unwinds double stranded nucleic acids using an ‘active’ or ‘passive’ mechanism has been a subject of much debate. Here, we derived analytic expressions for both the velocity and processivity of generic unwinding helicases, and showed that only by *simultaneously* using velocity and run-length data can the active/passive nature of a helicase, and hence the unwinding mechanism, be discerned. Simple expressions for the processivity or run-length of helicases were previously unavailable, hence our work should prove useful in the future analysis of helicase unwinding trajectories. Our results also quantitatively predict that the processivity should show a universal increase with force unlike the velocity, which has implications for *in vivo* unwinding of nucleic acids by the replisomal complex. This result seems to be borne out in a variety of experiments, and further work will help in verifying these remarkable predictions. Finally, we have also quantitatively shown for the T7 helicase, that the backstepping rate while unwinding is orders of magnitude larger than the backstepping rate while translocating on single-strand nucleic acid. This result is under-appreciated, and if taken into account could should qualitatively change the conclusions of the usual approaches of modeling, which assume that back stepping probabilities are negligible.

## Acknowledgments

This work was completed while SC was a graduate student at the Institute for Physical Sciences and Technology, University of Maryland. We acknowledge the National Science Foundation (CHE 16–36424 and DMR-1506969) and the Collie-Welch Foundation (F-0019) for supporting this work.

## APPENDIX

The full expressions for 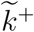, 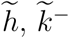 and 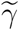 are given below:

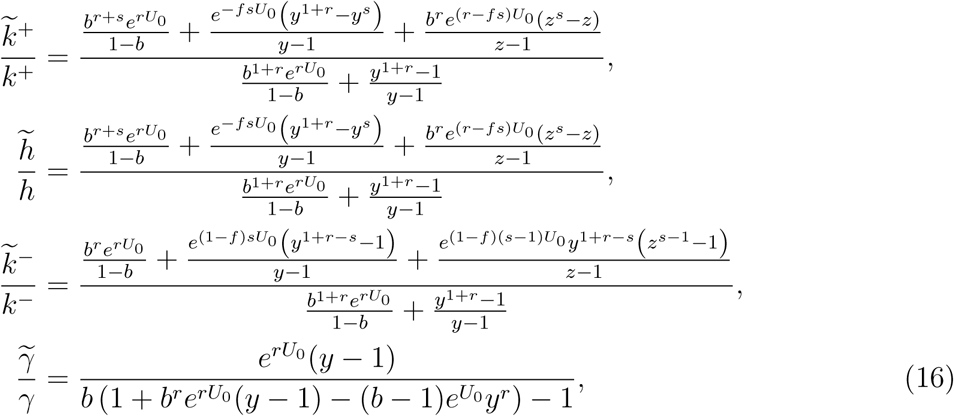
where 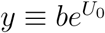 and 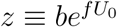.

